# Development and validation of glycolysis-cholesterol synthesis genes in lung adenocarcinoma

**DOI:** 10.1101/2022.08.15.503983

**Authors:** Bao Qian, Yangjie Guo, Jiuzhou Jiang

## Abstract

**Background:** Lung adenocarcinoma (LUAD) is the most common subtype of lung cancer which is one of the most dangerous malignant tumors affecting human health. The pathways of glycolysis and cholesterol production play an essential role in the metabolism of cancer. The mitochondrial pyruvate carrier (MPC) consists of MPC1 and MPC2, and impaired MPC function may induce a solid capacity for tumor proliferation, migration, and invasion.

**Methods:** Genes positively and negatively correlated with MPC1/2 expression were identified by calculating Spearman correlation, then gene ontology (GO) analysis was conducted. Univariate cox regression, lasso regression, and multivariate cox regression analyses were performed to model predictive outcome events using differentially expressed genes. Thirteen prognostic genes were selected to construct a prognostic model.

**Results:** 1359 and 2026 genes were positively and negatively associated with MPC1/2, respectively. The expression of MPC1 and MPC2 was significantly different. The prognostic model had great predictive performance in the test set.

**Conclusions:** MPC1/2 genes were involved in a cellular network associated with the malignant development of LUAD. The prognostic model can provide an essential basis for physicians to predict the clinical outcomes of LUAD patients.

## 1. Introduction

Although lung cancer is the second most prevalent cancer worldwide, it is the primary cause of cancer deaths, with an estimated 2 million new cases and 1.76 million deaths yearly [1, 2]. Fifteen percent were small cell lung cancer (SCLC), 85% were non-small cell lung cancer (NSCLC), and lung adenocarcinoma (LUAD) was the most common subtype of lung cancer [2–4].

Reprogramming of cell metabolism is an essential feature of malignancy, as shown by abnormal uptake of glucose and amino acids and dysregulation of glycolysis [5, 6]. Glycolysis is a specific metabolic pattern of tumor cells, which meets the requirements of tumor cells for ATP, etc.[7] Mitochondrial pyruvate carrier (MPC) consists of MPC1, and MPC2 is responsible for the import of pyruvate from the cytoplasmic matrix into the mitochondrial matrix, which can affect glycolysis, and damaged MPC function may induce tumors with solid capabilities for proliferation, migration, and invasion [8, 9]. Pyruvate is converted to acetyl coenzyme A, which is further changed to citric acid, a precursor substance required for lipogenesis, including the synthesis of cholesterol[10]. There is growing evidence for a close relationship between cholesterol metabolism and some types of cancer, such as allosteric interactions in the microenvironment of tumors, cancer cell spreading and metastasis forming, and lipid metabolism in tumor-initiating cells (TICs) [11, 12].

In recent decades, some studies have investigated potential prognostic signatures of LUAD using only bulk RNA-seq data, which mainly provides data on the average number of cells in the sample [13, 14]. Single-cell sequencing is a powerful instrument for dissecting the cellular and molecular landscape with single-cell resolution, revolutionizing our comprehension of the biological features and dynamics within cancer pathologies [15]. Single-cell RNA-seq technology can comprehensively characterize the heterogeneity of the tumor microenvironment and help dissect the complex cell type compositions and expressive heterogeneity in TME, and the Tumor Immune Single Cell Hub (TISCH) can assist us with a simple analysis. [16].

Our previous studies performed a series of analyses to select DEGs [17]. We analyzed the functions of genes that were positively or negatively associated with MPC1/2 and constructed a prognostic model. Then, we explored the correlation of the genes with the tumor microenvironment (TME). The risk-prognosis model can be used to formulate treatment options and analyze the prognosis.

## Method

### Analysis of DEGs

Based on our earlier research, gene expression matrices and clinical information of 585 LUAD patients were downloaded from The Cancer Genome Atlas (TCGA). These patients were divided into four groups based on co-expressed glycolysis and cholesterogenesis genes, and then survival analysis was conducted [17]. The two subtypes with the best and the worst prognosis were selected for differential expression analysis. 445 DEGs were identified using the “limma” package (V3.52.2), of which 221 were up-regulated, and 224 were down-regulated.

### Analysis of MPC1/2 Expression

By calculating Spearman correlation coefficients and corresponding false discovery rate (FDR) values (Beyer-Hardwick method) for MPC1/2 and other genes, we identified genes positively and negatively associated with MPC1/2 expression. After a series of filters, 1359 and 2026 genes were found to be positively and negatively correlated with MPC1/2 expression, respectively (Spearman correlation). To explore the possible functions and pathways of the identified positively and negatively associated genes, gene ontology (GO) analysis was conducted by applying the clusterProfiler package[18]. FDR < 0.05 was chosen as the criterion for cutoff.

### Prognostic gene selection and Construction and validation of a prognostic risk model

The TCGA-LUAD samples were randomly divided into training (50%) and test (50%) sets. Genes associated with prognosis were screened from DEGs using univariate Cox proportional hazard regression analysis based on clinical data of LUAD cases in TCGA (p < 0.01). We then perform Lasso regression using 10-fold cross-validation and a p-value of 0.05, the penalty parameter (λ) is determined by the minimum criterion. Thirteen prognostic genes were selected to construct a prognostic model. We explored the relationship between prognostic genes and tumor microenvironment based on Tumor Immune Single Cell Hub Database (TISCH).

A prognostic risk model based on prognostic gene expression was constructed using multivariate Cox regression analysis. Then, we calculated the risk score for each sample as follows: 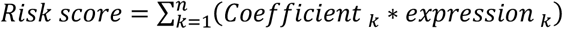. According to the median risk scores, the samples were divided into high and low-risk groups. A training set and a test set were used to validate the model’s robustness and plot the risk score distribution as well as the ROC curve and KM survival curve.

## Result

### MPC complexes as possible regulators of the glycolysis-cholesterol synthesis axis in tumors

Among the four metabolic subtypes, the expression of MPC1 and MPC2 was significantly different (Figure 1A). The mean levels of MPC1 and MPC2 expression were higher in patients in the cholesterogenic group than in the other subtypes, and MPC1 expression was significantly higher in the cholesterogenic group than in the mixed group in particular (FDR <0.05). Previous studies have shown that MPC1 is absent or underexpressed in various cancers and is associated with poor prognosis[19]. To explore the pathways correlated with MPC1/2 expression levels, we conducted a correlation analysis between MPC1/2 and all the other genes. In total, 1359 and 2026 genes were found to be positively and negatively associated with MPC1/2, respectively (Spearman correlation, Figure 1B). Further GO functional enrichment analysis showed that genes with a positive association with MPC1/2 were implicated in the mitochondrial inner membrane, mitochondrial, protein-containing complex, mitochondrial matrix, and ribosome (Figure 1C); genes with a negative association with MPC1/2 were associated with histone modification, ameboidal-type cell migration, cell junction assembly, cell-substrate adhesion, and external encapsulating (Figure 1D). MPC1/2 genes are engaged in a cellular network associated with the malignant development of LUAD.

**Figure 1.**
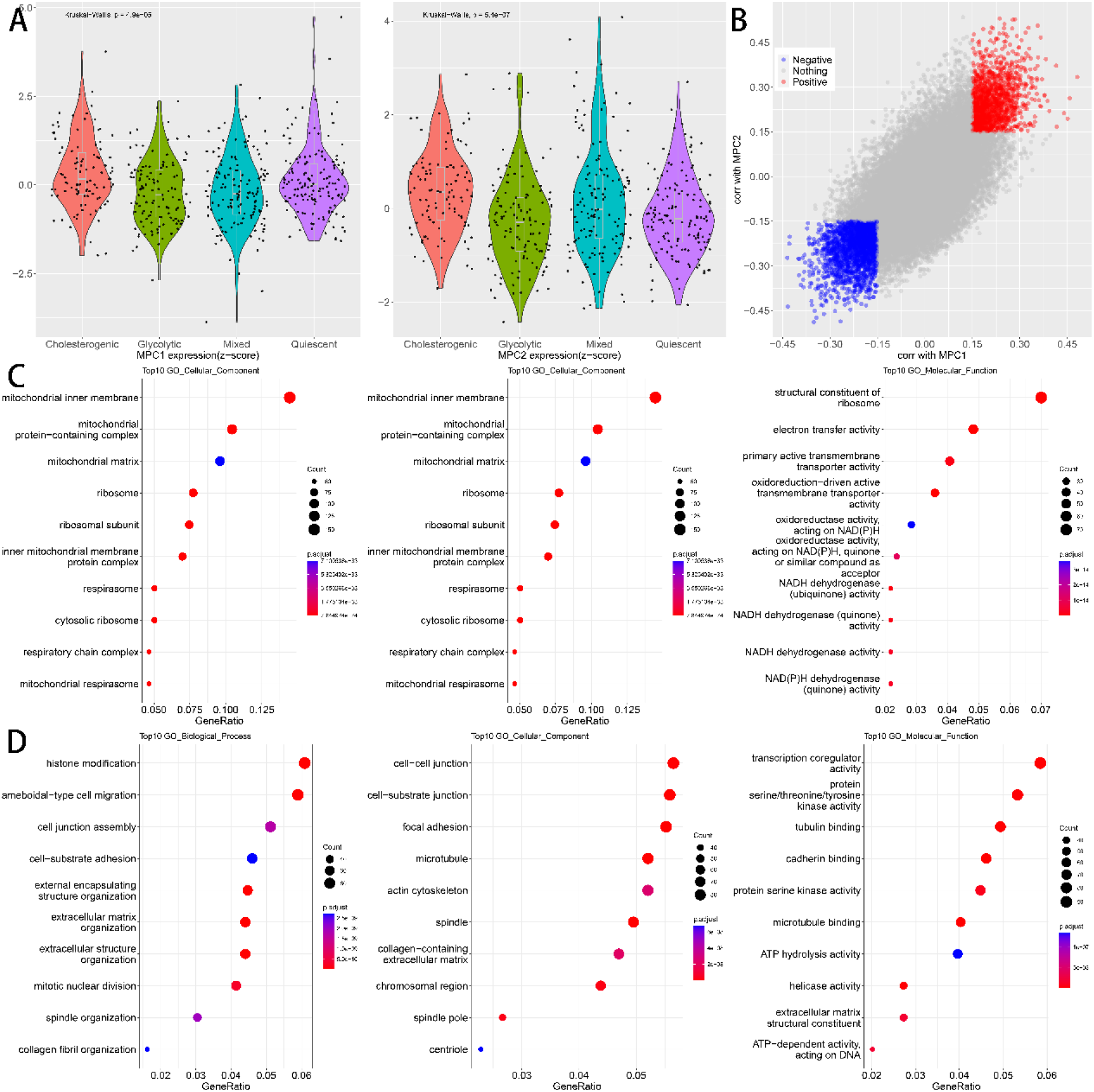
(A) MPC1/MPC2 expression across different metabolic subtypes. (B) Scatter plot describing the correlation between MPC1 (X-axis) and MPC2 (Y-axis). (C) Results of GO enrichment analysis of genes positively associated with MPC1/2. (D) Results of GO enrichment analysis of genes negatively associated with MPC1/2.

### Identification and assessment of 13 prognostic genes for LUAD

Based on the survival dataset of LUAD patients, a univariate Cox regression was applied to the expression profiles of 445 DEGs. A total of 199 DEGs were identified according to the p < 0.01 criterion. We conducted Lasso regression on these genes and identified 13 differentially expressed genes associated with LUAD prognosis (Figure 2A). The optimum value of the penalty parameter was identified through 10 rounds of cross-validation (Figure 2B). By multiple Cox regression analysis,11 prognostic genes, including CTSV, UBE2SP1, RP11-627G23.1, COL4A3, ADAMTS7P3, COL6A6, MIR4697HG, PKP2, IGF2BP1, MIR31HG, KLK11, SERPIND1, HHIPL2 were selected to construct the prognostic model (Figure 2C). These 13 genes were then used to calculate a risk score for each sample based on their expression levels. Risk score = - 0.046*CTSV + 0.452*UBE2SP1-0.064*RP11-627G23.1+0.016*COL4A3-0.172*ADAMTS7P3 - 0.103*COL6A6 - 0.030*MIR4697HG + 0.146*PKP2 + 0.014*IGF2BP1 + 0.375*MIR31HG - 0.071 *KLK11 + 0.052*SERPIND1 + 0.026*HHIPL2.

**Figure 2.**
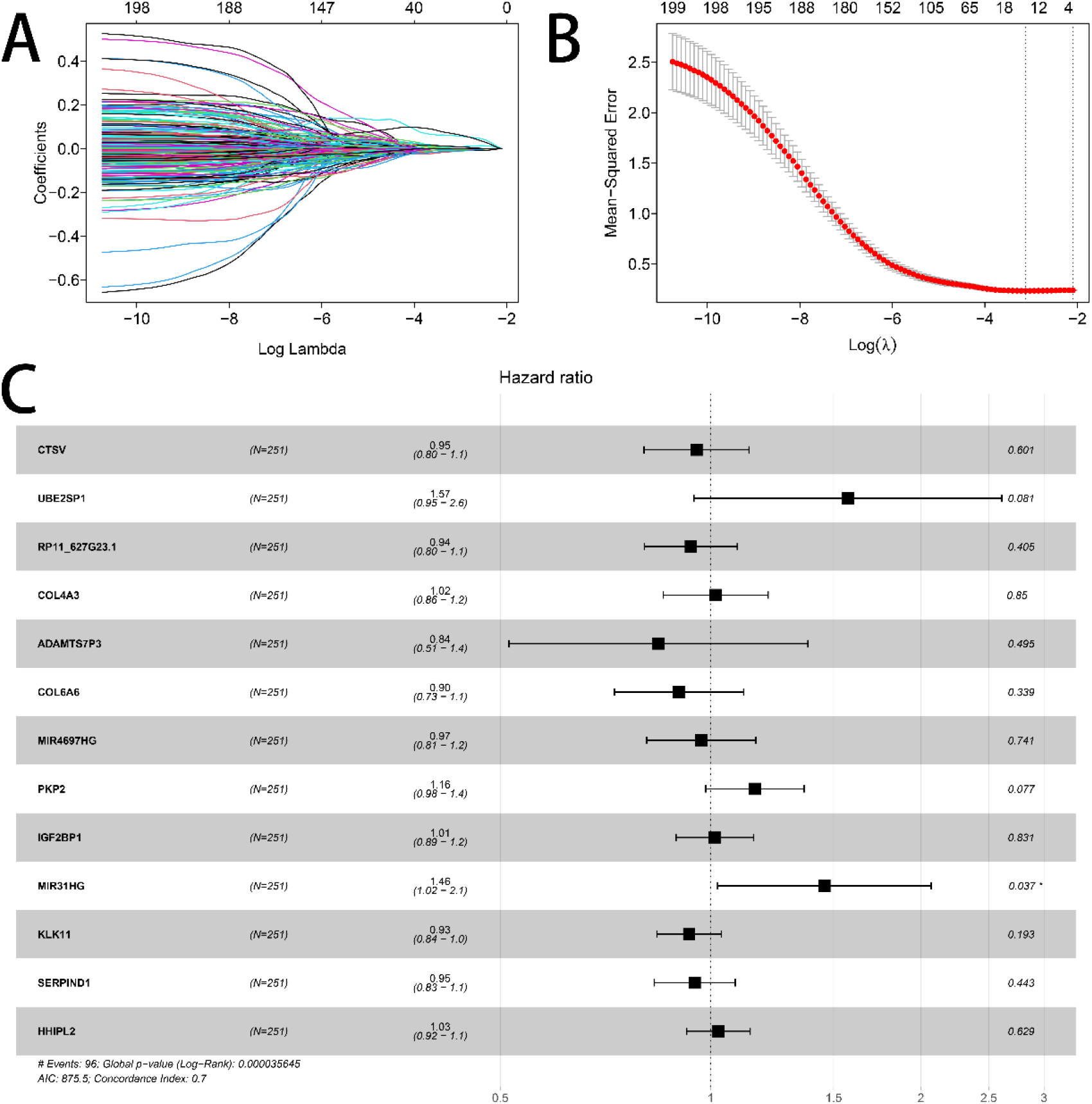
(A) LASSO regression of the 13 prognostic genes. (B) Determination of optimal values of penalty parameters by 10 rounds of cross-validation. (C) Forest plot for multivariate cox regression.

### Model evaluation

Use each sample’s 13 prognostic gene expression levels to calculate and plot the risk score distribution and the gene expression heat map for the training set (Figure 3A). Survival times were significantly shorter in the high-risk score samples than in the low-risk score samples, indicating that samples with lower risk scores were more likely to have a better prognosis. There were 126 samples with risk scores greater than the median classified as the high-risk group and 125 samples below the median classified as the low-risk group. There was a significant prognostic difference between the high-risk and low-risk groups (P<0.05, Figure 3B). Figure 3C shows the performance of the predictive classification based on risk scores at 1, 3, and 5 years in the training set.

**Figure 3.**
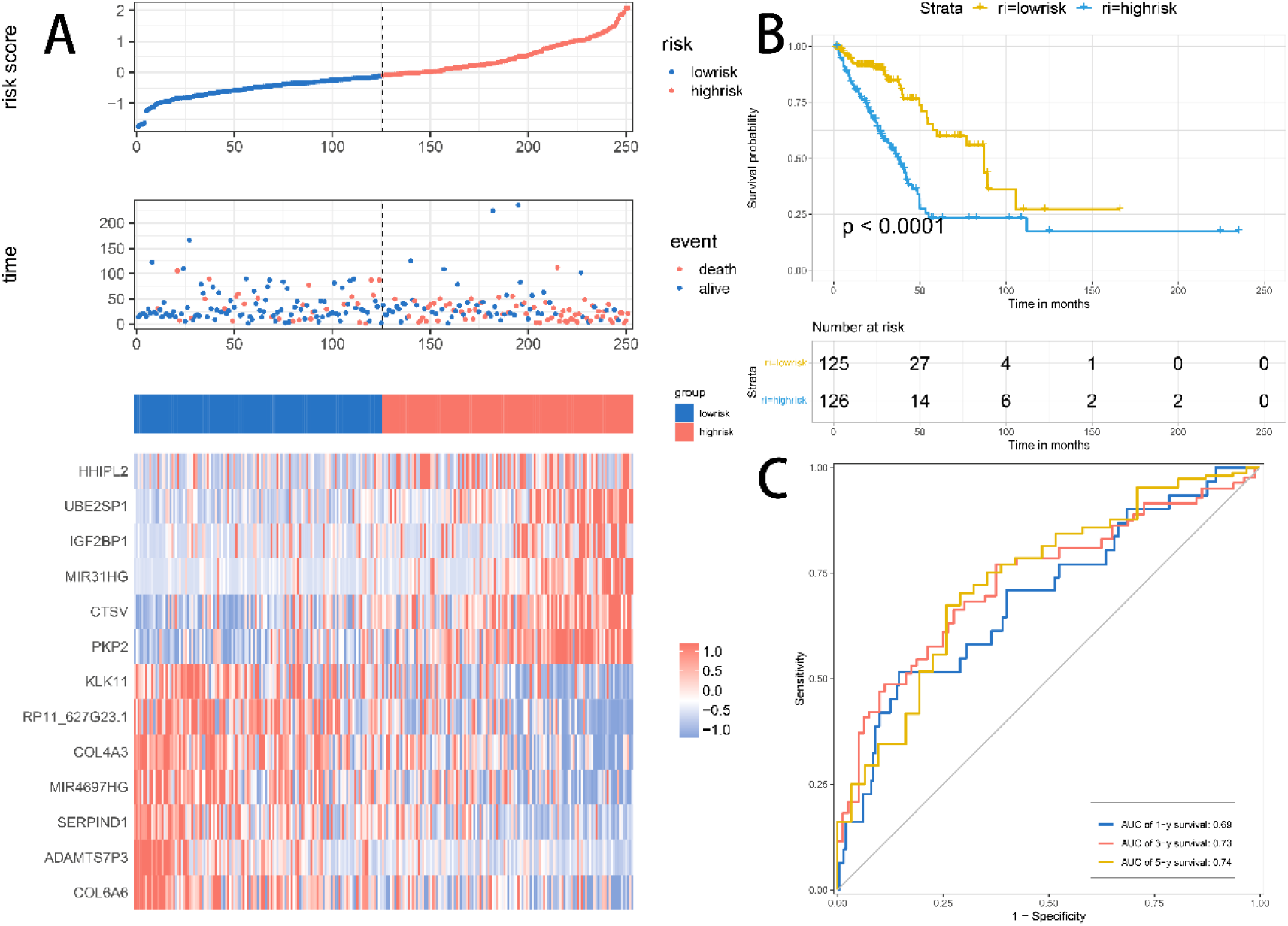
(A) Distribution of risk scores, survival time, and heatmap of expression of 9 genes in the training set. (B) The KM survival curve of the risk model in the training set. (C) The ROC curve of the risk model in the training set.

Risk score distributions and gene expression heatmap of the test set were shown in Figure 4A. Consistent with the trend in the training set, indicating a poorer prognosis for samples with high-risk scores. The KM survival curve showed that there was a significant prognostic difference between the high-risk and low-risk groups in the test set (Figure 4B). Figure 4C shows the performance of the predictive classification based on risk scores at 1, 3, and 5 years in the test set.

**Figure 4.**
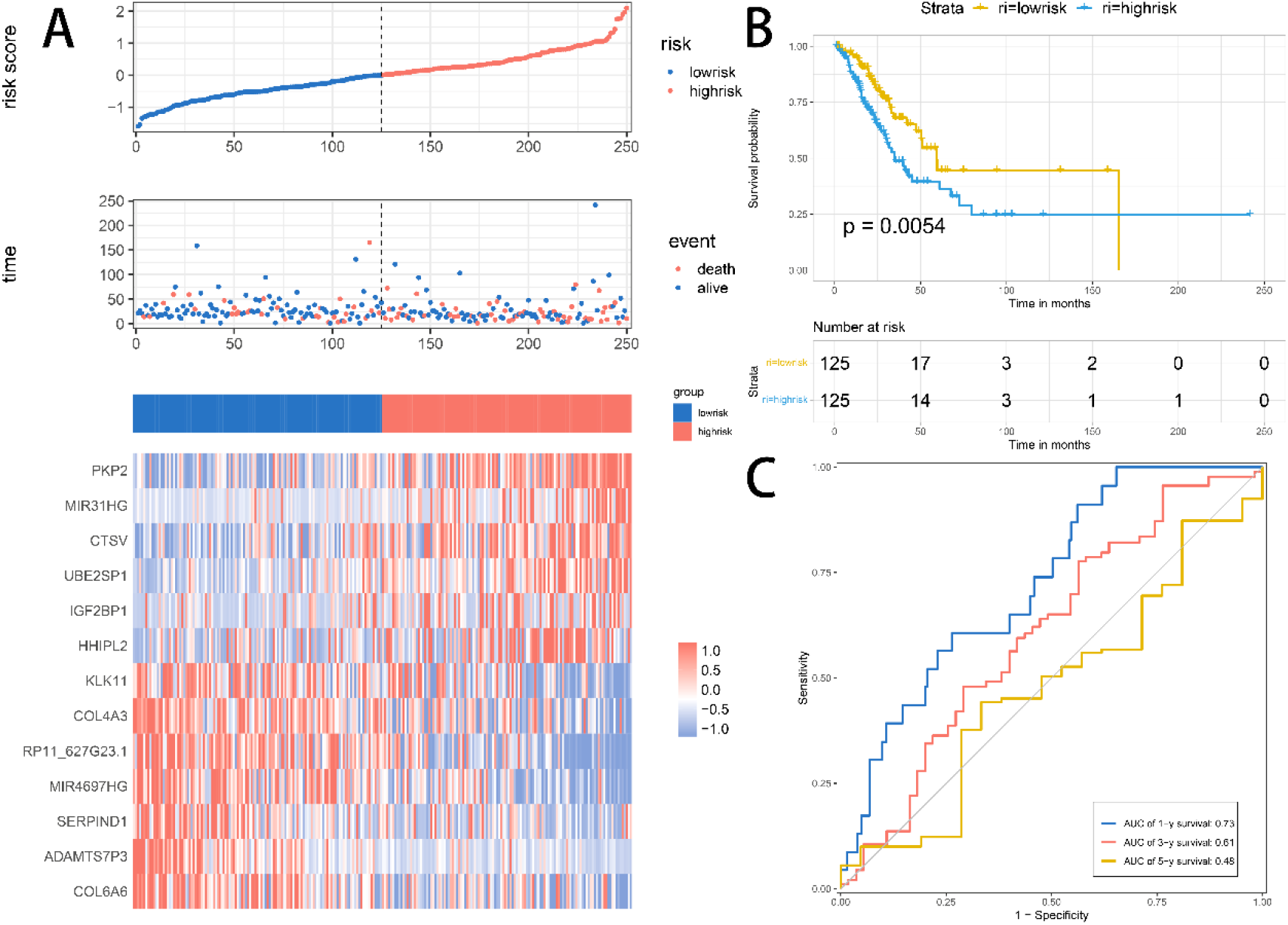
(A) Distribution of risk scores, survival time, and heatmap of expression of 9 genes in the test set. (B) The KM survival curve of the risk model in the test set. (C) The ROC curve of the risk model in the test set.

### Analysis of the correlation between prognostic genes and TME

Given the role of TME in tumor development and its impact on prognosis, we used a NSCLC_GSE117570 dataset from the TISCH database to analyze the expression of some prognostic genes in TME-associated cells. We then examined the dataset, which is categorized into ten cell types. Figure 5A shows the number of cells of each cell type and presents the distribution of each type of TME-associated cells. In this dataset, malignant cells were the most abundant (n = 2721). We found that CTSV, COL4A3, KLK11, and PKP2 had higher expression in malignant cells compared to other types of TME-associated cells (Figure 5B). These results support the association of prognostic genes with lung cancer.

**Figure 5.**
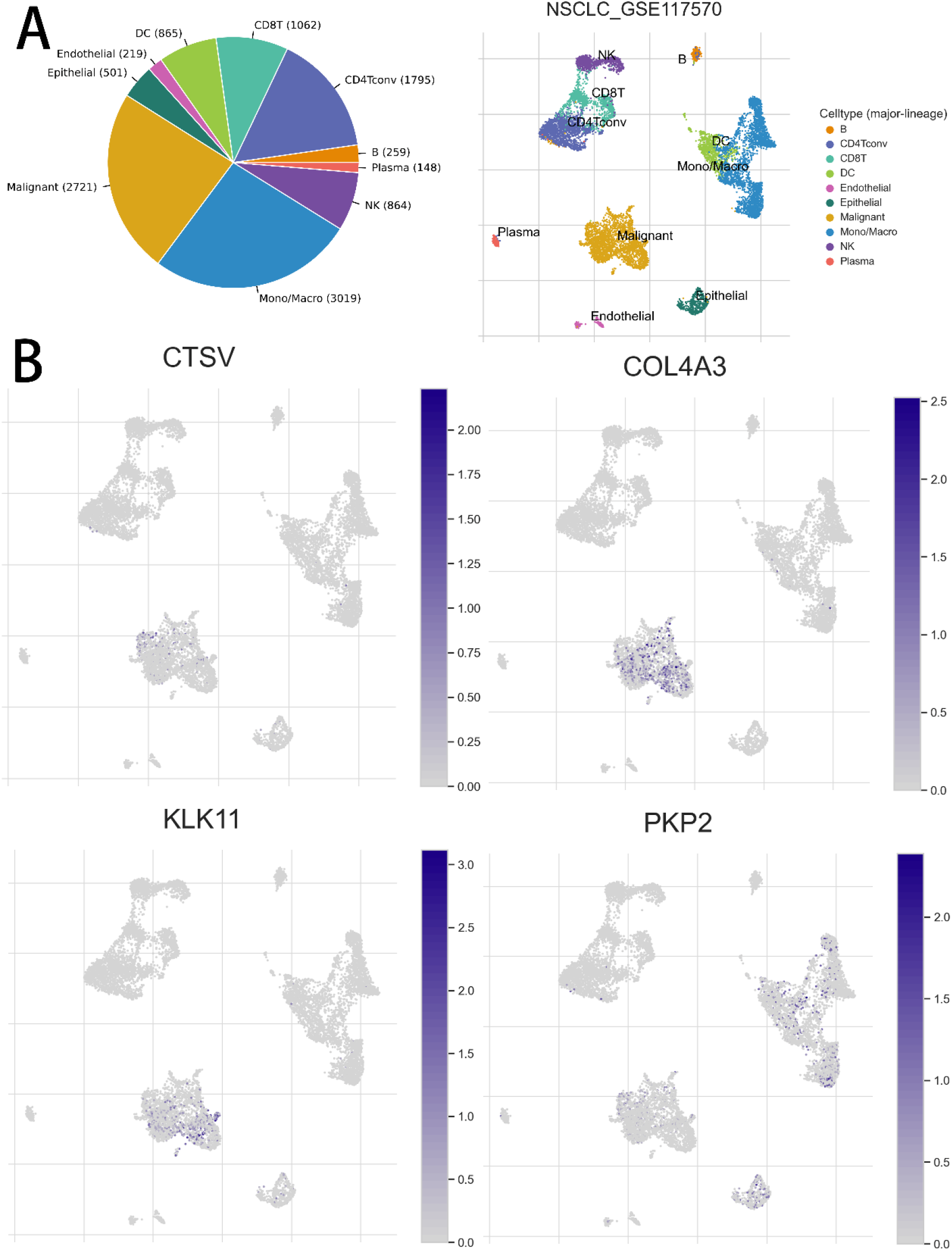
(A) Number of cells of each cell type in the NSCLC_GSE117570 dataset, with a description of the distribution of TME-associated cells of each type. (B) Distribution of CTSV, COL4A3, KLK11, and PKP2 in TME-associated cell types.

## Discussion

The majority of patients with advanced lung cancer, one of the most deadly malignancies in humans, are recurrent and treatment resistant. The glycolysis and cholesterol production pathways play an essential role in cancer metabolism. Therefore, it is essential to clarify the metabolic pathways of lung cancer for its prevention and treatment.

Pyruvate is central to carbohydrate, fat, and amino acid metabolism. Pyruvate is appealing as a therapeutic target against cancer because it promotes respiratory reserve capacity and mitochondrial oxygen consumption, which may contribute to the aggressive disease phenotype [20]. Mitochondrial pyruvate carrier(MPC) is one of the critical enzymes responsible for pyruvate transport and oxidation [9]. Low or absent MPC1 and MPC2 levels lead to metabolic disorders and alterations in tumor metabolism, and their restored expression inhibits tumor growth, invasiveness, metastasis, and stemness[21]. By analyzing the expression of MPC1/2, the results showed that there were significantly different expressions of MPC1/2 among different subtypes of metabolism, suggesting that the MPC complex affects the metabolic pathway and thus participates in the malignant progression of lung cancer by regulating the amount of pyruvate entering the mitochondria.

Based on 93 glycolysis and cholesterol synthesis genes, patients were assigned to four subtypes: glycolytic, cholesterogenic, quiescent, and mixed. Cholesterol subtypes have a better prognosis than other subtypes. We performed a survival analysis and identified DEGs between the best and worst prognosis subtypes. Nine prognostic genes were selected. Reviewing the literature, we found that the role of many prognostic genes has been studied concerning lung cancer and has been revealed to impact tumorigenesis and progression. CSTV has been previously shown to be a promising biomarker for lung cancer by Yang et al. [22]. Similarly, COL6A6, IGF2BP1, and MIR31HG have been found to play important roles in lung cancer proliferation and may be potential therapeutic targets [23–25]. Analyzing the association between these genes and TME, we discovered that some prognostic genes were highly expressed in malignant cells using a single-cell sequencing database, which contributed to our construction of a better prognostic model. The model was robust on the training and test datasets and had a great predictive performance.

This study also has some limitations. First, there are no external tests of our model. Secondly, future experimental verification is needed.

## Conclusions

MPC1/2 genes were involved in a cellular network associated with the malignant development of LUAD. The prognostic model can provide an important basis for physicians to predict the clinical outcomes of LUAD patients.

